# Real-time TIRF imaging of single adiponectin vesicle exocytosis in 3T3-L1 adipocytes stably expressing mCherry fused to human adiponectin

**DOI:** 10.1101/2022.09.13.507762

**Authors:** Man Mohan Shrestha, Sebastian Barg, Charlotta S. Olofsson

## Abstract

Adiponectin is a peptide hormone abundantly released from adipocytes, and reduced circulating levels are associated with obesity-related diseases, such as type 2 diabetes, insulin resistance and cardiovascular disease. Adiponectin is released by regulated exocytosis of secretory vesicles, but traditional molecular biology and imaging techniques lack the specificity and time resolution to adequately quantify exocytosis and trafficking of adiponectin-containing vesicles. Here we generated 3T3-L1 adipocytes that stably express mCherry-tagged human adiponectin, resulting in robust labelling of small adiponectin vesicles with a diameter of 200-300 nm, in live cells. Total internal reflection fluorescence (TIRF) microscopy was used to visualise and quantify exocytosis and adiponectin release in real-time, observed as rapid disappearance of the fluorescence of individual vesicles. Bulk adiponectin secretion measurements confirmed that the labelled adiponectin was secreted to the surrounding solution under these conditions, and expressed in the same vesicle population as endogenous adiponectin. In contrast to previous electrophysiological results, elevation of cytosolic Ca^2+^ alone was sufficient to induce exocytosis, although at a lower rate compared to elevated cytosolic cAMP. We conclude that the adiponectin-mCherry-labelled cells are useful for studying adiponectin exocytosis at the single vesicle level, and that an intracellular elevation of either cAMP or Ca^2+^ can trigger adiponectin vesicle release.

## Introduction

The adipocyte-derived hormone adiponectin has anti-diabetic, anti-atherogenic and anti-inflammatory effects and acts as an insulin sensitizer. Thus, adiponectin plays crucial roles in metabolic disorders, including insulin resistance, type 2 diabetes and cardiovascular disease (1, 2). Overexpression of adiponectin is associated with protection against both acute and chronic effects of high fat diet-induced lipotoxic effects of lipid accumulation, and enhances the metabolic flexibility of adipose tissue (3).

A number of studies support that adiponectin is secreted via regulated exocytosis of pre-formed adiponectin-containing vesicles. Lim et al. demonstrated the vesicular localisation of adiponectin and that insulin stimulates release of measurable amounts of adiponectin at time-points between 60 and 120 minutes (4). The insulin-induced secretion was unaffected by pre-incubation with the protein synthesis inhibitor cyclohexamide, indicating that the secretagogue stimulates release of vesicles belonging to a pool of already synthesised vesicle. Our own work has defined an alternate pathway for adiponectin exocytosis: By combining electrophysiological and biochemical techniques with mathematical modelling, we have demonstrated that adiponectin exocytosis is triggered by an increase of intracellular cAMP and activation of exchange factor directly activated by cAMP, isoform 1 (Epac1). The cAMP/Epac1-dependent adiponectin release occurs in the absolute absence of Ca^2+^ (shown by buffering with the Ca^2+^ chelator BAPTA), but is none the less potentiated by an elevation of cytosolic Ca^2+^. This regulation of adiponectin exocytosis exists in cultured 3T3-L1 adipocytes as well as in primary mouse and human adipocytes. Membrane capacitance recordings (that measure exocytosis as the increase in plasma membrane area that occurs upon vesicle fusion) showed that the cAMP/Ca^2+^-triggered adiponectin exocytosis can be detected within 1-2 min after intracellular addition of stimuli. The stimuli consisted of inclusion of cAMP with or without Ca^2+^ in the patch pipette solution washing into the recorded cell, or extracellular addition of the cAMP-elevating agent forskolin and/or the Ca^2+^ ionophore ionomycin (5–7). Later work demonstrated that pre-treatment of adipocytes with cyclohexamide was without effect also on the cAMP-triggered adiponectin secretion (8). Collectively, our work proposes that pre-stored adiponectin vesicles undergo rapid exocytosis upon stimulation with cAMP in both the presence and absence of Ca^2+^, in agreement with the cell model in Fig. 1.

**Fig. 1.**
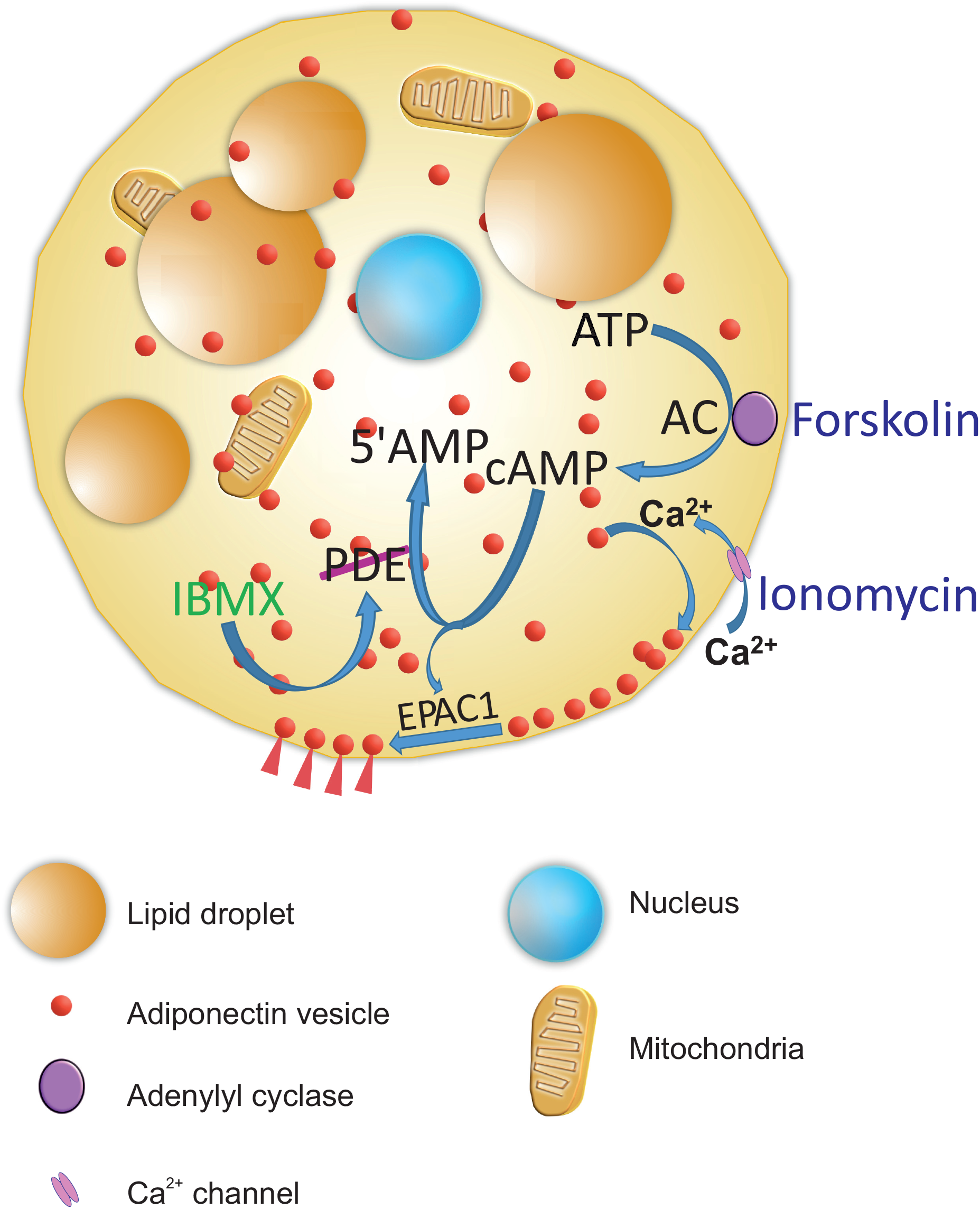
Effects of an elevation of cAMP and Ca^2+^ on adiponectin exocytosis in white adipocytes. Forskolin and 3-isobutyl-1-methylxanthine (IBMX) elevate the intracellular concentration of cAMP through the activation of adenylyl cyclase (AC) and inhibition of phosphodiesterase, respectively. The elevation of cAMP results in activation of the downstream target Epac1 and fusion of adiponectin containing vesicles with the plasma membrane with subsequent adiponectin release. Ionomycin, a calcium ionophore, facilitates the transport of Ca^2+^ across the plasma membrane and the elevation in intracellular Ca^2+^ augments the fusion of adiponectin-containing vesicles with the plasma membrane. Ca^2+^ also induces replenishment of release-competent adiponectin vesicles, to maintain secretion of adiponectin over longer time periods.

Total internal reflection fluorescence (TIRF) microscopy is an optical technique that enables the excitation of fluorophores in a very narrow optical section. TIRF microscopy provides a means to selectively excite fluorophores near the adherent cell surface, while counter-minimizing fluorescent signals from other intracellular regions (10). The narrow TIRF excitation field allows high-resolution visualisation of processes occurring near or in the plasma membrane, in live cells (11). Thus, by applying TIRF microscopy to adipocytes expressing a fluorophore localised to adiponectin vesicles, their exocytosis can be visually observed at single cell and single vesicle level, at high spatial and temporal resolution.

Here, we describe the generation of 3T3-L1 cells that stably express mCherry fused to the C-terminal end of human adiponectin, to produce a genetically homogenous clonal cell population that over-express the recombinant protein of interest. We demonstrate that the adiponectin-mCherry expressing cells can be differentiated into mature adipocytes, as judged by lipid-filling. Using TIRF microscopy, we can visualise and quantify Ca^2+^ and cAMP-triggered adiponectin exocytosis in real-time, at the single-vesicle resolution.

## Material and methods

### Safety and Handling of Recombinant Lentiviruses

All the lentiviruses were generated using a third generation packaging system as per Biosafety Level 2 and institutional guidelines of the University of Gothenburg. Disposable plastic pipettes were used and liquid waste was decontaminated with 10 % bleach. Laboratory materials in contact with viral particles were treated as biohazardous waste and autoclaved.

### Cell culture

HEK293T (ATCC) cells were cultured in high glucose DMEM (Invitrogen, GIBCO) supplemented with 10 % fetal bovine serum (FBS) and 1 % penicillin-streptomycin at 37 ^0^C and 5 % CO_2_. Cultured clonal 3T3-L1 fibroblasts (Zenbio) were seeded and differentiated into mature adipocytes and used for experiments at day 8 or 9 according to previous establish protocol (7). In brief, 3T3-L1 fibroblasts were cultured in high glucose DMEM (Invitrogen, GIBCO) supplemented with 10 % new born calf serum at 37 ^0^C and 5 % CO_2_. Two days after confluence, cells were washed twice with warm PBS and a differentiation medium containing 10 % FBS, 1 % penicillin-streptomycin, 0.85 μM insulin, 0.5 mM 3-isobutyl-1-methylxanhine and 1 μM dexamethasone was added for 2 days. The differentiation medium was there after replaced with medium containing 10 % FBS and insulin for another 2 days. Cells were then maintained in DMEM with 10 % FBS. Adipocytes were used for experiments on day 8 or 9 post differentiation. Only 3T3-L1 adipocytes that appeared mature, as judged visually by accumulated lipid, were studied.

### Generation of lentiviral constructs

Three candidate hairpin sequences for mouse adiponectin (NM_009605.5) each containing 21 sense bases that are identical to the target gene, a loop containing an XhoI restriction site and 21 antisense bases that are complementary to the sense bases followed by a poly T termination sequence, were designed using an small interfering RNA (siRNA) selection tool hosted by the Whitehead Institute for Biomedical Research (http://sirna.wi.mit.edu/) (12). Starting at 25nt downstream of the start codon (ATG), we searched for 21nt sequences that matched the pattern AA(N19). It was ascertained that the G-C content was >35 %. To minimize degradation of off-target mRNAs, the NCBI’s BLAST program was used. Since some sequences are likely more effective than others, multiple target sequences were selected for the *Acrp30* gene (Table 1).

**Table 1:**
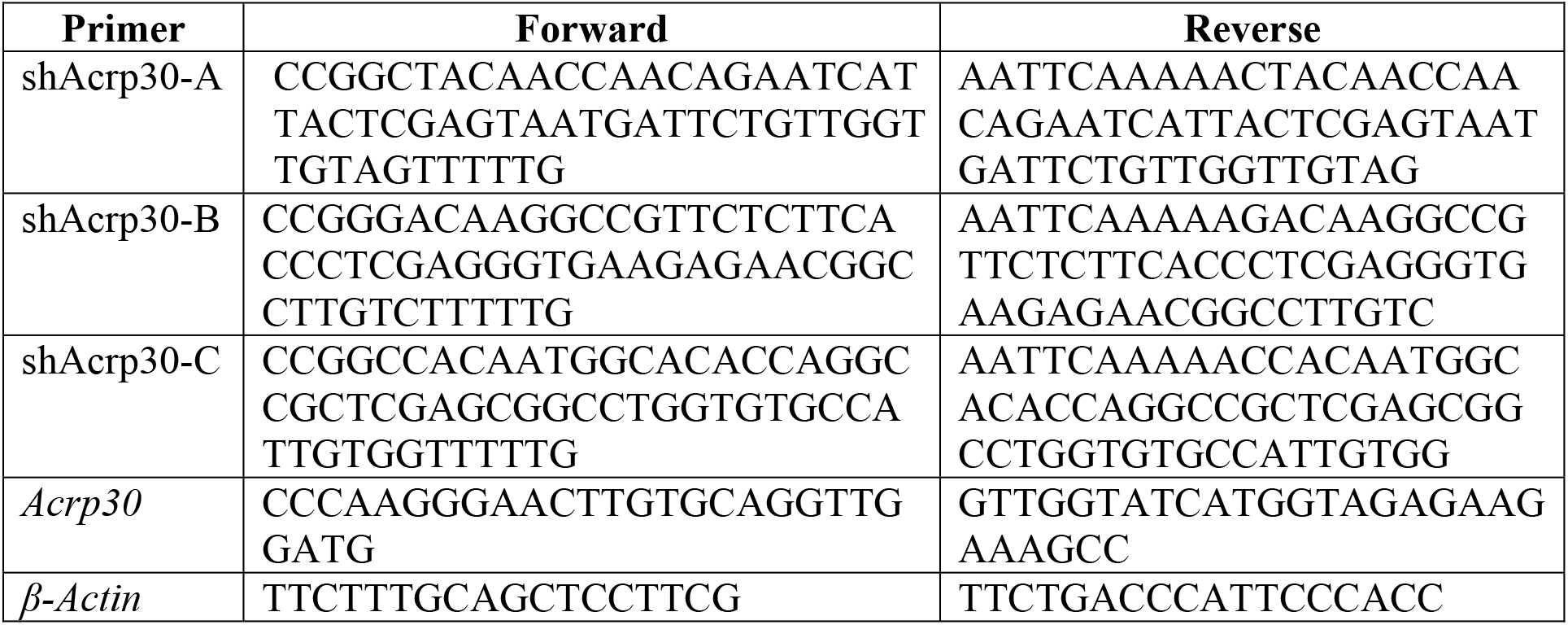

The annealed oligos were inserted into an empty backbone third generation transfer plasmid (pLKO.1-TRC cloning vector; Plasmid #10878; addgene) using AgeI (5’ cloning site) and EcoRI (3’ cloning site) restriction digestion sites. Similarly, a candidate hairpin sequence for green fluorescent protein (GFP) was used for a control/scramble (pLKO.1 GFP; Plasmid #30323; addgene). mCherry fused to human adiponectin (hAcrp30-mCherry) was cloned into a pLenti vector using NheI and HpaI restriction digestion sites to generate lentiviral constructs (Supplementary Fig. 1A). The sequences of the knockdown and lentiviral constructs were verified using restriction digestion and Sanger’s Sequencing from Eurofins Genomics.

### Lentivirus Packaging and Infection

Lentiviruses were produced by co-transfection of HEK293T cells with lentiviral expression and packaging plasmids using the calcium phosphate transfection method (13, 14). Viral supernatant was collected after 48 hours of transfection, centrifuged at 1,000 x g for 5 minutes and filtered through 0.45 µm filter. The viruses were further concentrated by ultracentrifugation using Ti45 rotor in Beckman Coulter Optima-L-100 XP at 116,000 x g for 2 hours at 4 °C. For gene silencing, 3T3-L1 fibroblasts were transduced with titrated viruses in presence of 8 µg/mL of polybrene (Hexadimethrine bromide, Cat. No: 107689-10G, Sigma-ALDRICH) and subjected to antibiotic selection using 4 µg/mL puromycin after 48 hours. The successful stable expression was verified by fluorescence microscopy for fusion proteins whereas knockdown efficiency was assessed by Western blotting and RT-qPCR at protein and mRNA level respectively. For generation of a stably expressing mCherry fused adiponectin cell line, 3T3-L1 fibroblasts were infected with optimum amount of viral particles in the presence of polybrene. The growth medium was changed 24 hours after infection.

### Adiponectin secretion

3T3-L1 cells stably expressing adiponectin-mCherry were seeded and differentiated into mature adipocytes on 35 mm glass bottom dish (D35-20-1.5-N, Cellvis). The mature adipocytes were incubated with an extracellular solution containing 140 mM NaCL, 3.6 mM KCL, 0.5 mM MgSO_4_, 0.5 mM NaH_2_PO_4_, 2.0 mM NaHCO_3_ and 5.0 mM Hepes (pH 7.4) supplemented with 2.6 mM CaCl_2_ and 5.0 mM glucose for 30 minutes at 32 °C. Test substances were added for 30 min at 32 °C under gentle shaking. At termination of the incubation, the medium containing secreted product was collected, and the adipocytes were lysed using PBS containing 2 % SDS and protease inhibitor (1 tablet per 10 mL; cOmplete Mini, Roche Diagnotics). Secretion samples were centrifuged for 5 minutes at 2000 rpm and 4 ^0^C, and the supernatant was stored at -80 °C. Secreted HMW and total adiponectin (measured with HMW and Total Adiponectin ELISA (human) kit, 47-ADPMS-E01, Alpco) were expressed in relation to total protein content. Cell lysates were centrifuged at 14000 rpm for 10 minutes at 4 ^0^C and total protein contents were measured by Bradford protein assay.

### Lipolysis Assays

Matured adipocytes were incubated with an extracellular (EC) solution for 30 minutes at 32 °C. Lipolysis was stimulated by adding the adenylate cyclase activator Forskolin (10 µM) and the phosphodiesterase inhibitor isobutylmethylxanthine (IBMX) (200 µM) or Ionomycin (1 µM) in serum-free extracellular (EC) solution for 30 min at 32 °C under gentle shaking. At termination of the incubation, the medium containing secreted product was collected, and the adipocytes were lysed using PBS containing 2% SDS and protease inhibitor (1 tablet per 10 mL; cOmplete Mini, Roche Diagnotics). Cell lysates were centrifuged at 14000 rpm for 10 minutes at 4 ^0^C and total protein contents were measured by Bradford protein assay. Secretion samples were centrifuged for 5 minutes at 2000 rpm and 4 ^0^C, and the supernatant was collected. Lipolysis was assessed from the release of FFA and glycerol in the secretion samples using non-esterified fatty acids measurement kit (FUJIFILM Wako Chemicals Europe GmbH) and free glycerol reagents (G7793 and F6428, Sigma) respectively and normalised to total protein contents.

### Real-Time Quantitative PCR

RNA was extracted and purified using QIAzol (Qiagen) and ReliaPrep RNA Cell MiniPrep System (Promega). 1 µg of RNA was used for reverse transcription. cDNA synthesis was performed according to the protocol as described in Invitrogen^TM^ SuperScript^TM^ II Reverse Transcriptase kit (Thermo Fisher Scientific #18064022) in presence of 40 units of RNaseOUT^TM^ Recombinant Ribonuclease Inhibitor (Thermo Fisher Scientific #10777-019) and 200 units of Reverse Transcriptase per 20 µL of reaction mixture. The real-time quantitative PCR (qPCR) assays were performed using Fast SYBR Master Mix (4385612, Applied Biosystems). The relative expression of mRNA abundance was calculated using 2^-ΔΔCt^ and was normalized to β-actin. Data are represented as mean ± SEM, ****p<0.0001 (n=3).

### Western blotting

3T3-L1 adipocytes were lysed with TNET buffer (50 mM Tris-HCl, pH 7.5, 150 mM NaCl, 10 mM NaF, 1 mM EDTA and 1% Triton X-100) containing 2 mM Na_3_VO_4_ (Sodium Orthovanadate) and 1 mM pmsf (phenylmethylsulfonyl fluoride) and 1x protease inhibitor cocktail tablet (Roche). Whole cell lysates were centrifuged at 12,000 rpm at 4 ^0^C for 10 minutes. The protein concentration was determined using Bradford protein assay. Proteins from whole cell lysate (20 µg) were separated by 8 % sodium dodecyl sulfate polyacrylamide gel electrophoresis (SDS/PAGE) and then transferred onto polyvinylidene difluoride (PVDF) membranes (Millipore) at 120 V for 2 hours using transfer buffer (25 mM Tris-base, pH 8.3, 190 mM glycine, 20 % methanol and 0.1 % SDS). After blocking in blocking buffer (2 % BSA in PBS containing 0.2 % Tween-20 and 0.01 % sodium azide) for 1 hour at room temperature, the membrane was incubated overnight at 4 ^0^C with primary antibody against adiponectin (ab3455, Abcam) and tubulin (T9026, Sigma) as loading control at 1:1000 dilution. The primary antibodies were probed using HRP-conjugated secondary antibodies at 1:5000 dilution and detection was performed with enhanced chemiluminescence.

### Live Cell imaging using TIRF microscopy and analysis of vesicle exocytosis

3T3-L1 adipocytes expressing hAcrp30-mCherry were pre-treated with extracellular solution for 30 minutes at 32 °C and kept in the same solution supplemented with 2.6 mM CaCl_2_ and 5.0 mM glucose during imaging. Image acquisition was performed by TIRF Observer Z1 microscope with alpha Plan-Apochromat 100×/1.46 Oil objective (both Carl Zeiss) and an EMCCD camera (Evolve 512 delta, Photometrics). Images were acquired using Zeiss Zen 2.6 (blue edition). Excitation laser of wavelength 561 nm with exposure time 25–100 ms was used for mCherry. Forskolin together with the phosphodiesterase inhibitor 3-isobutyl-1-methylxanthine (FSK/IBMX) or ionomycin alone were added 1–2.5 minutes after acquisition start and the time point was noted. TIRF images were acquired over 15 minutes at 2 Hz through 1801 total readings. Circles of region of interest (ROI) around the vesicles were used to measure the fluorescence intensities of images over time. All the readings were normalized to the intensity measured at time point zero, after removal of the background fluorescence. Values were plotted against time, so as to indicate the time course of adiponectin containing vesicle exocytosis. Vesicle size was estimated using fluorescent polystyrene beads of known diameter. Vesicle locations were found by eye, a 21 pxl long line was centered on this location, and the brightness along this line was read out (“linescan”). The following piecewise linear function describing a trapezoid shape was then fitted to the linescan to estimate spot width: if (x1< x<x1+bl) y=a+s*(x-x1); if (x2-bl< x<x2) y=a+s*(x2-x); if (x1+bl<x<(x2-bl) y=a+s*bl; else y=a, where x1 and x2 are boundaries of the peak, a the peak amplitude, s the slope, bl the width of the two sloped sections, and y the brightness. The apparent width is w=x2-x1-bl, which is the full width at half maximum (FWHM) of the trapezoid. bl was fixed to 1.5pxl (240nm), the theoretical resolution of the system. Beads with known diameters were imaged and measured in the same way, to create a reference scale.

### Analysis

Images were analysed using Zeiss Zen 2.6 (blue edition) and OriginPro (OriginLab Corporation). All data are presented as mean values ± SEM for designated number of experiments. Significance of difference was tested by Student’s t-test and ANOVA. Statistical analysis were performed using Prism 9.0 (GraphPad, San Diego, CA).

## Results

### Knock down and rescue of endogenous adiponectin in 3T3-L1 adipocytes

3T3-L1 cells with low passage number were used and it was ascertained that the cells were able to express high levels of endogenous adiponectin when differentiated into adipocytes (Fig. 2A). To avoid that native adiponectin interferes with the expression of human adiponectin-mCherry, the endogenous adiponectin was knocked down by short hairpin RNA (shRNA). Three different target sequences for shRNA mouse adiponectin knockdown were designed: shAcrp30-A, shAcrp30-B and shAcrp30-C (Table 1). The targets were inserted into a lentiviral plasmid and 3T3-L1 fibroblasts were transduced with the viral particles. The transduced cells were thereafter differentiated into mature adipocytes. For two out of three knockdowns (shAcrp30-A and shAcrp30-C) transduced cells could be cultured into mature adipocytes, whereas cells transduced with shAcrp30-B did not survive. The successful knockdown was confirmed by protein and gene expression, where adiponectin gene knockdown was compared with scrambled shRNA targeting GFP (Fig. 2B-D).

**Fig. 2.**
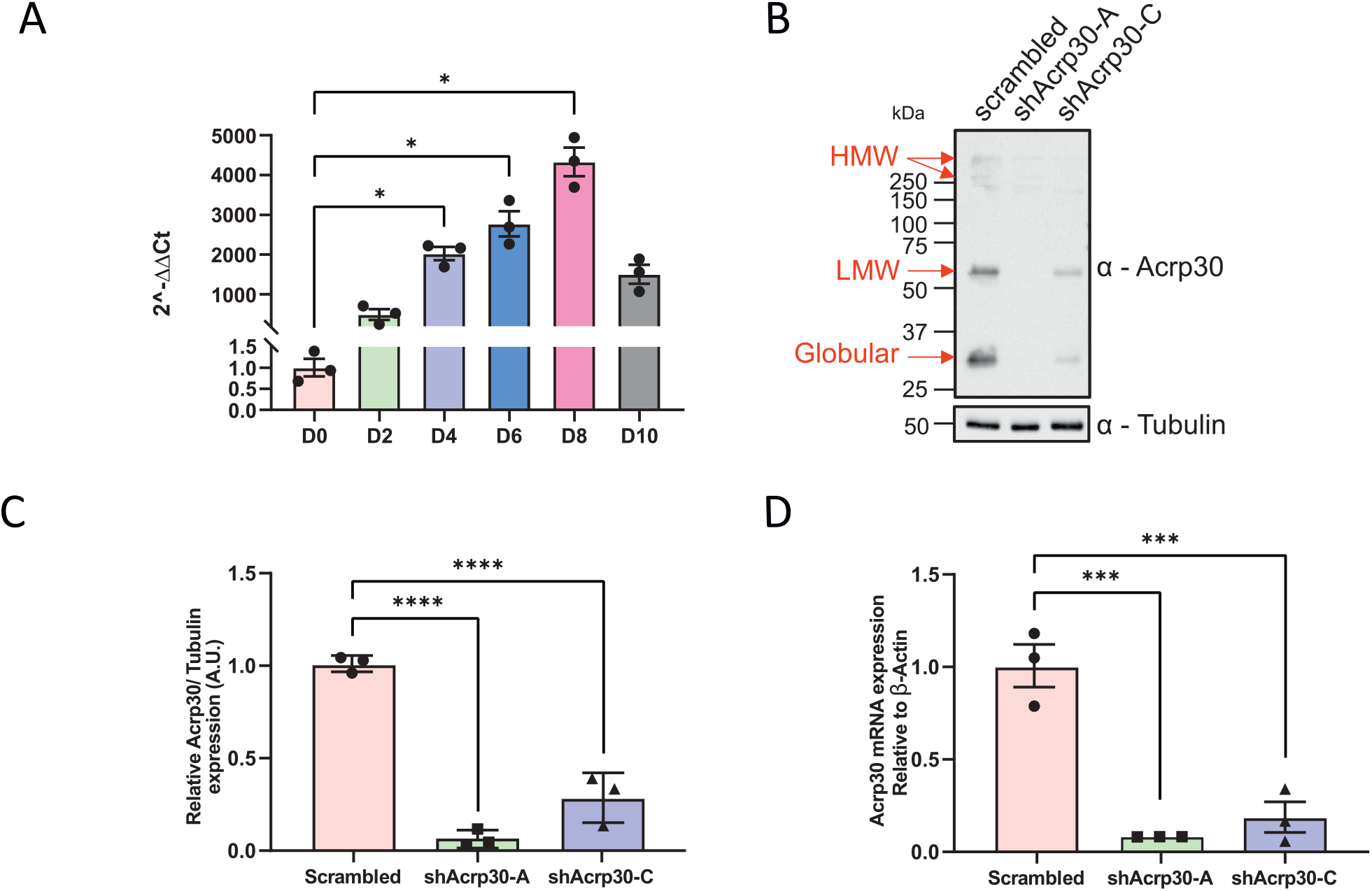
Expression of adiponectin and shRNA-mediated adiponectin knockdown in 3T3-L1 adipocytes. **A)** Gene expression of endogenous adiponectin in 3T3-L1 cells/adipocytes measured at every 48 hours after initiation of differentiation until day 10. Western blot **(B**) and densitometric analysis (**C**) of adiponectin protein expression after shRNA-mediated knockdown of endogenous adiponectin in 3T3-L1 cells. The cells were infected with lenti-virus-expressing control shRNA targeting GFP (scrambled) and two variants of shRNA targeting murine adiponectin (shAcrp30-A and shAcrp30-C), and differentiated into mature adipocytes after puromycin selection. **D)** mRNA levels of adiponectin in shRNA-mediated knockdown compared with control cells. Statistical analysis: Brown-Forsythe and Welch ANOVA tests (Dunnett’s T3 multiple comparisons test), assuming unequal SDs at α – level of 0.05 for A and Ordinary One-way ANOVA tests (Dunnett’s multiple comparisons test), assuming equal SDs at α – level of 0.05 for C & D. Data are presented as mean ± SEM of three independent experiments. *P<0.05, ***P<0.001 and ****P<0.0001.

Cells that displayed the most efficient knockdown (shAcrp30-A), were rescued with hAcrp30-mCherry alone (plasmid map in Suppl. Fig. 1) or with a FLAG tag either at the N- or C-terminal in the fibroblast stage. (The FLAG was added to allow future protein purification by affinity chromatography as well as localisation of the fusion protein in live cells.) The rescued cells were seeded and differentiated into mature adipocytes, and the over-expression of hAcrp30 was verified by Western blotting using antibodies against adiponectin or FLAG. The hAcrp30-mCherry tagged with FLAG at the C-terminal was found to be highly over-expressed compared to the other two variants under both reduced (Fig. 3A) and non-reduced (Fig. 3B) conditions. The over-expression of adiponectin-mCherry was visualized by inverted fluorescence microscope (Fig. 3C) and TIRF microscopy (Fig. 3D); TIRF imaging demonstrated a punctate pattern of fluorescence, as typical for vesicular compartments. Most of the labelled vesicles appeared diffraction limited or slightly larger in size. Comparing the apparent width (FWHM) of the vesicles with that of fluorescently labelled beads of known diameter allowed us to estimate their median diameter to ∼200-300 nm (Fig. 3E, F).

**Fig. 3.**
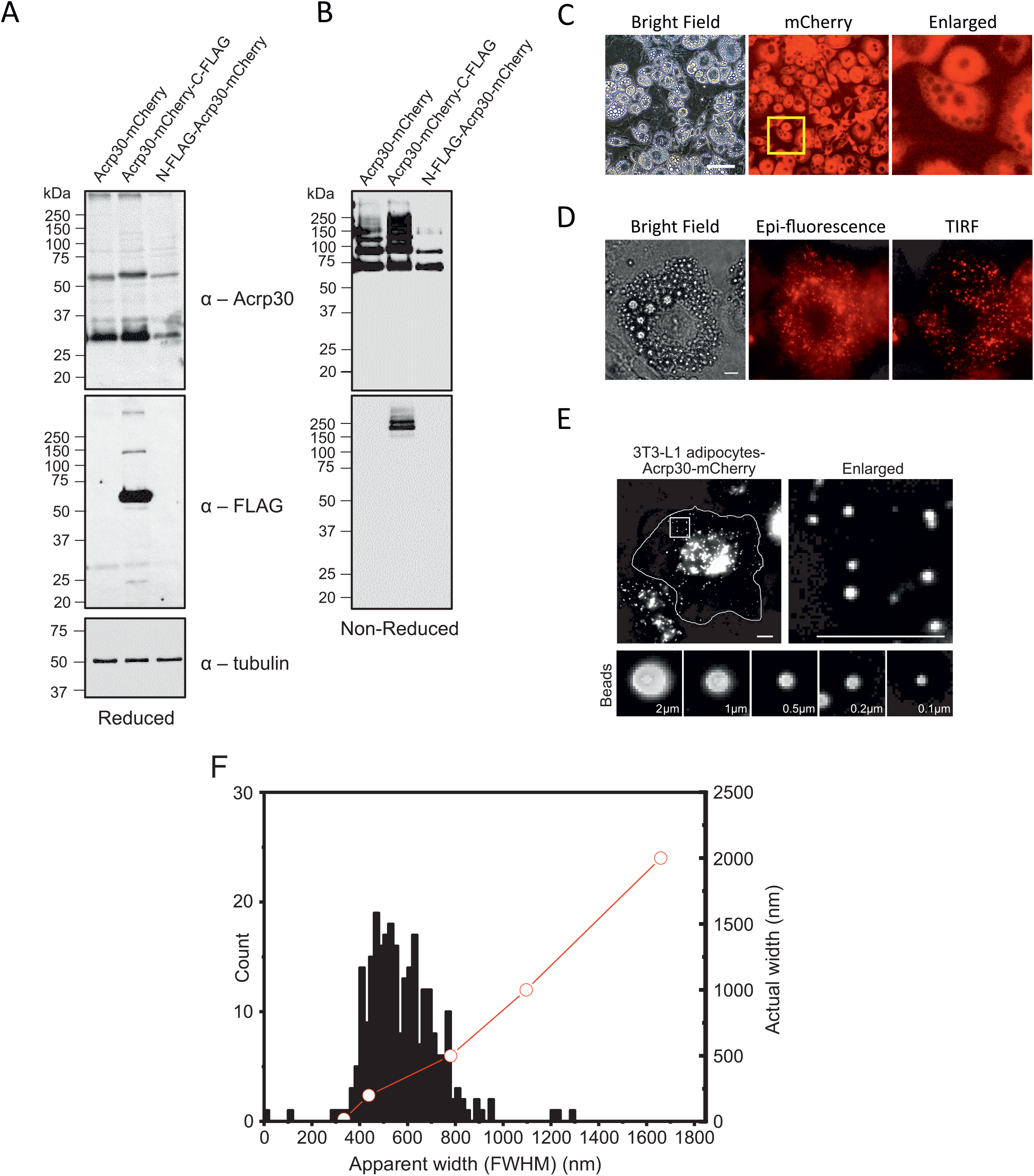
Rescue of shRNA-mediated adiponectin knockdown with mCherry fused with human adiponectin and estimation of vesicle size. Western blot for **A)** reduced and **B)** non-reduced adiponectin. The expression of mCherry-fused adiponectin in the 3T3-L1 adipocytes was observed under **C)** Fluorescence (Scale bar=100 µm) and **D)** TIRF microscope (Scale bar=10 µm). The 3T3-L1 cells, with shRNA-mediated silencing of endogenous adiponectin expression, were infected with lenti-virus for stably expressing human adiponectin fused with mCherry at C-terminal alone or with FLAG-tag at either C- or N-terminal. The lenti-virus infected cells were differentiated and the over-expression of recombinant adiponectin was analysed. **E-F)** Estimation of adiponectin-mCherry vesicle size (n=267), as described in Methods. Fluorescent polystyrene beads with a diameters of 2, 1, 0.5, 0.2 and 0.1 μm were imaged. Scale bars=10 μm.

### An elevation of intracellular cAMP or Ca^2+^ triggers exocytosis of mCherry-labelled adiponectin vesicles

We have shown that a combination of the adenylyl cyclase activator forskolin and the phosphodiesterase inhibitor 3-isobutyl-1-methylxanthine (FSK/IBMX) increases 3T3-L1 adipocyte cAMP levels >6-fold (7), and that the Ca^2+^ ionophore ionomycin elevates the intracellular Ca^2+^ level ([Ca^2+^]_i_; (6). Thus, we investigated adiponectin vesicle exocytosis in differentiated 3T3-L1 adipocytes stably expressing adiponectin-mCherry exposed to FSK/IBMX or ionomycin. As shown in Fig. 4A, addition of FSK/IBMX led to disappearance of mCherry-labelled vesicles (bottom images in Fig. 4A and Suppl. video 1). A rapid loss of fluorescence puncta was observed upon stimulation, which we interpreted as exocytosis events (top row Fig. 4C and Fig. 4D). Approx. 60% of the vesicles observed by TIRF microscopy fused with the plasma membrane after addition of the stimulatory agents, while the remaining 40% were stable throughout the recording (middle row Fig. 4C and Fig. 4E). This is expected since it is unlikely that all adiponectin vesicles that reside close to the plasma membrane, within the TIRF zone, are primed for exocytosis (15). In experiments with adipocytes exposed to the control solution during the entire experiment (Fig. 4B and Suppl. video 2), the labelled adiponectin vesicles remained stable throughout the TIRF recording (bottom row Fig. 4C). Calculation of the fraction of released vesicles confirmed that a larger number of vesicles were exocytosed upon exposure to the cAMP-elevating agents compared to control (Fig. 4F). At the end of 15 min recordings, significantly fewer adiponectin-mCherry labelled vesicles remained in adipocytes exposed to FSK/IBMX, compared to control solution (Fig. 4G).

**Fig. 4.**
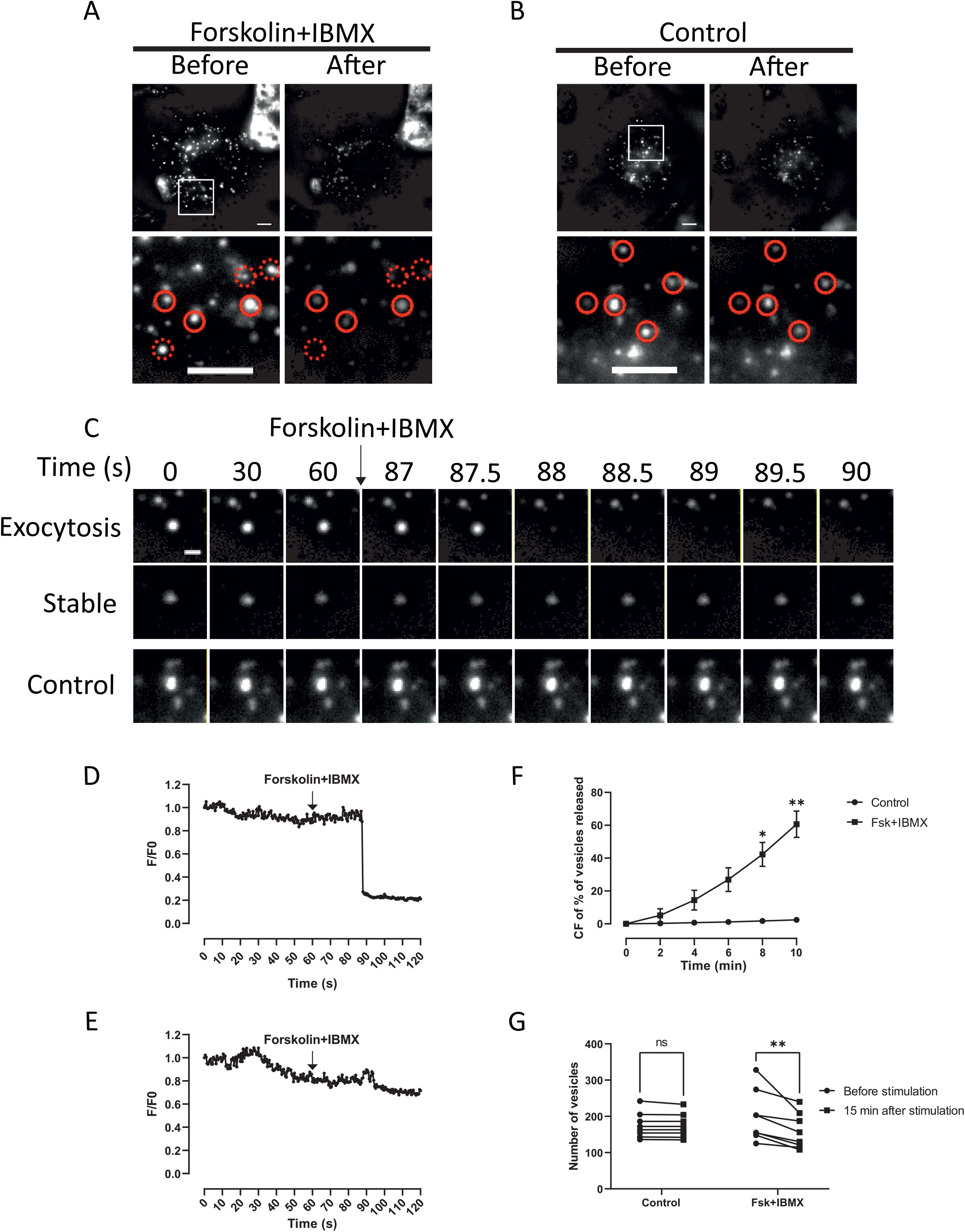
TIRF imaging of cAMP-stimulated 3T3-L1 adipocyte adiponectin exocytosis. 3T3-L1 adipocytes stably expressing recombinant adiponectin were serum starved for 30 min at 32^0^C and treated with **A**) 10 µM Forskolin and 200 µM IBMX or **B)** DMSO (Control) for 15 minutes during which images were taken at 2 FPS using TIRF microscopy. A laser wavelength of 561 nm with exposure time between 30 - 100 ms was used to excite mCherry. Note in inset region at lower panel that some vesicles are released (dotted circles) while others remain stable (whole circles). Scale bar = 10 µm. **C)** Examples of FSK/IBMX-induced single vesicle dynamics over time, in the stimulated adipocyte in (A) and in the control adipocyte in (B). Scale bar = 2 µm. **D-E)** Relative average fluorescence intensity change vs. time in a single vesicle stimulated with FSK/IBMX. **F)** Cumulative frequency of percentage of vesicles released at the end of a 10 min FSK/IBMX stimulation. **G)** Vesicle count before and 15 min after stimulation with FSK/IBMX. Statistical analysis: 2way ANOVA (Sidak’s multiple comparisons test). Data are presented as mean ± SEM of eight independent experiments. *P<0.05; **P<0.01.

Ionomycin likewise triggered the exocytosis of adiponectin-containing vesicles (Fig. 5A, and Suppl. video 3). Similar to the experiments with FSK/IBMX, a rapid loss of fluorescence puncta was observed upon stimulation, indicating vesicle exocytosis (top row Fig. 5C and Fig. 5D) and a fraction of adiponectin-containing vesicles remained stable throughout the recording (middle row Fig. 5C and Fig. 5E). It was again confirmed that adiponectin-mCherry vesicles remained stable in adipocytes maintained in control solution (Fig. 5B, bottom row Fig. 5C and Suppl. video 4). The fraction of released vesicles was significantly larger in the ionomycin-exposed adipocytes compared to control experiments, at all investigated time-points (Fig. 5F). Significantly fewer adiponectin-mCherry labelled vesicles remained in ionomycin-exposed adipocytes compared to control cells, at the end of 15 minute recordings (Fig. 5G). As can be seen upon comparison of Figs. 4F and 5F, whereas 60% of the vesicles residing in the TIRF zone were released after 10 min exposure to FSK/IBMX, the corresponding fraction of secreted vesicles amounted to only 15% in ionomycin-treated adipocytes.

**Fig. 5.**
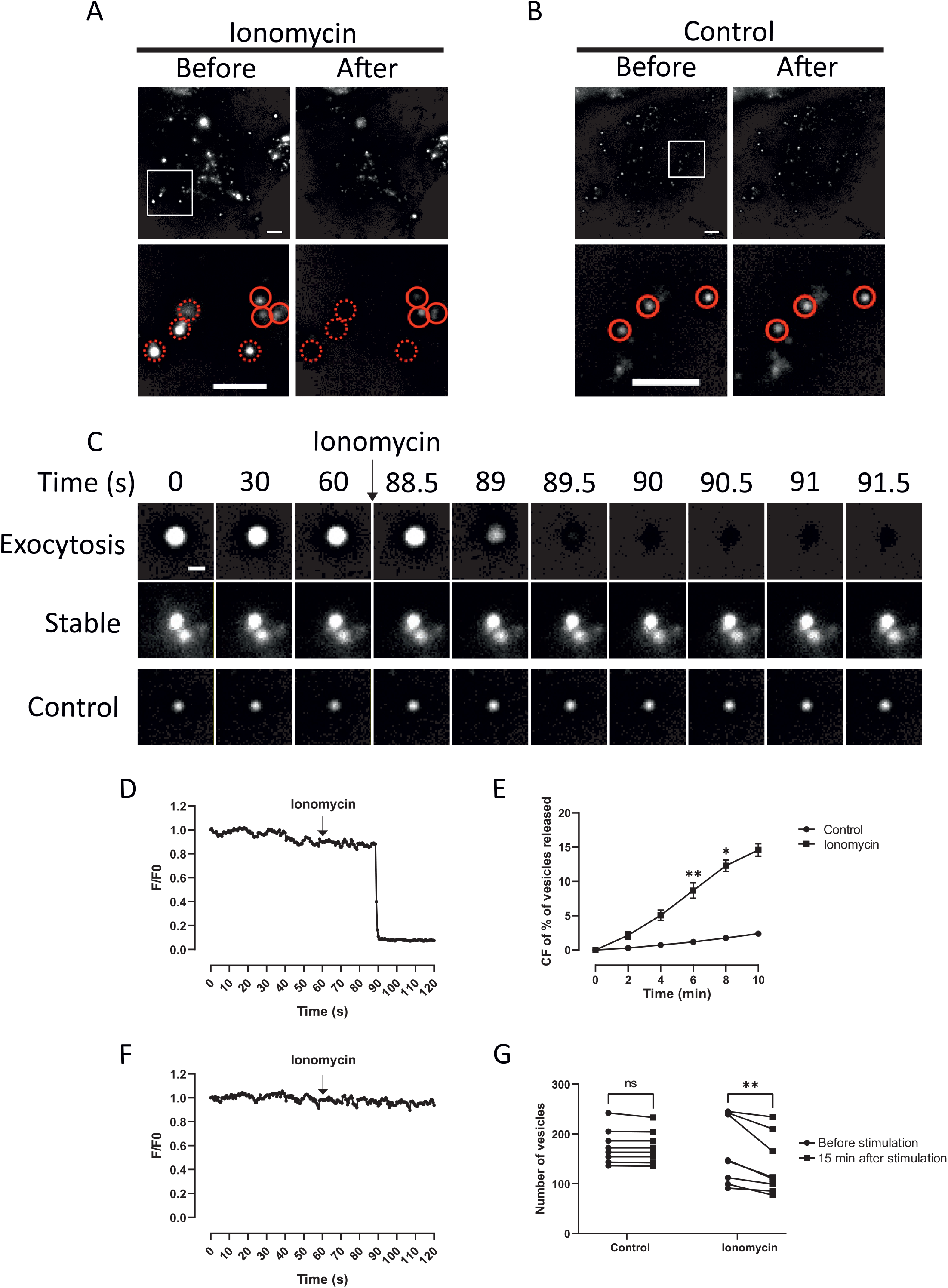
TIRF imaging of Ca2+-stimulated 3T3-L1 adipocyte adiponectin exocytosis. 3T3-L1 adipocytes stably expressing recombinant adiponectin were serum starved for 30 min at 32^0^C and treated with **A)** 1 µM ionomycin or **B)** DMSO (Control) for 15 minutes during which images were taken at 2 FPS using TIRF microscopy. A laser wavelength of 561 nm with exposure time between 30 - 100 ms was used to excite mCherry. Note in inset region at lower panel that some vesicles are released (dotted circles) while others remain stable (whole circles). Scale bar = 10 µm. **C)** Examples of ionomycin-induced single vesicle dynamics over time, in the stimulated adipocyte in (A) and in the control cell in (B). Scale bar = 2 µm. **D-E)** Relative average fluorescence intensity change vs. time in a single vesicle upon ionomycin stimulation. **F)** Cumulative frequency of percentage of vesicles released by ionomycin at the end of 10 min of stimulation. **G)** Vesicles count before and 15 min after stimulation with ionomycin. Statistical analysis: 2way ANOVA (Sidak’s multiple comparisons test). Data are presented as mean ± SEM of eight independent experiments. *P<0.05; **P<0.01.

### Measurements of released adiponectin, FFA and glycerol in 3T3-L1 adipocytes stably expressing adiponectin-mCherry

To verify the adiponectin secretion capacity of the stably expressing 3T3-L1 cells, the matured adipocytes were incubated in the presence of FSK/IBMX or ionomycin and product secreted to the medium was measured by ELISA. In agreement with previous results (6, 7) the secretagogues stimulated the release of mouse adiponectin in native 3T3-L1 adipocytes (Fig. 6A). The 3T3-L1 adipocytes stably expressing adiponectin-mCherry likewise secreted adiponectin in response to both FSK/IBMX and ionomycin and both total (Fig. 6B) and the high-molecular weight (HMW) form (Fig. 6C) of human adiponectin were released over basal (unstimulated control cells). The data confirm that the stably expressing adipocytes secrete smaller and larger forms of the adiponectin-mCherry fusion protein in response to cAMP or Ca^2+^ stimulation, at levels similar to that observed in mouse native adipocytes. To further validate the functionality of in the adiponectin-mCherry adipocytes, we measured release of free fatty acids (FFA) and glycerol. FSK/IBMX elevated levels of both FFA and glycerol in the incubation medium whereas ionomycin was without effect of release of either of the lipid metabolites (Figs. 6D, E).

**Fig. 6.**
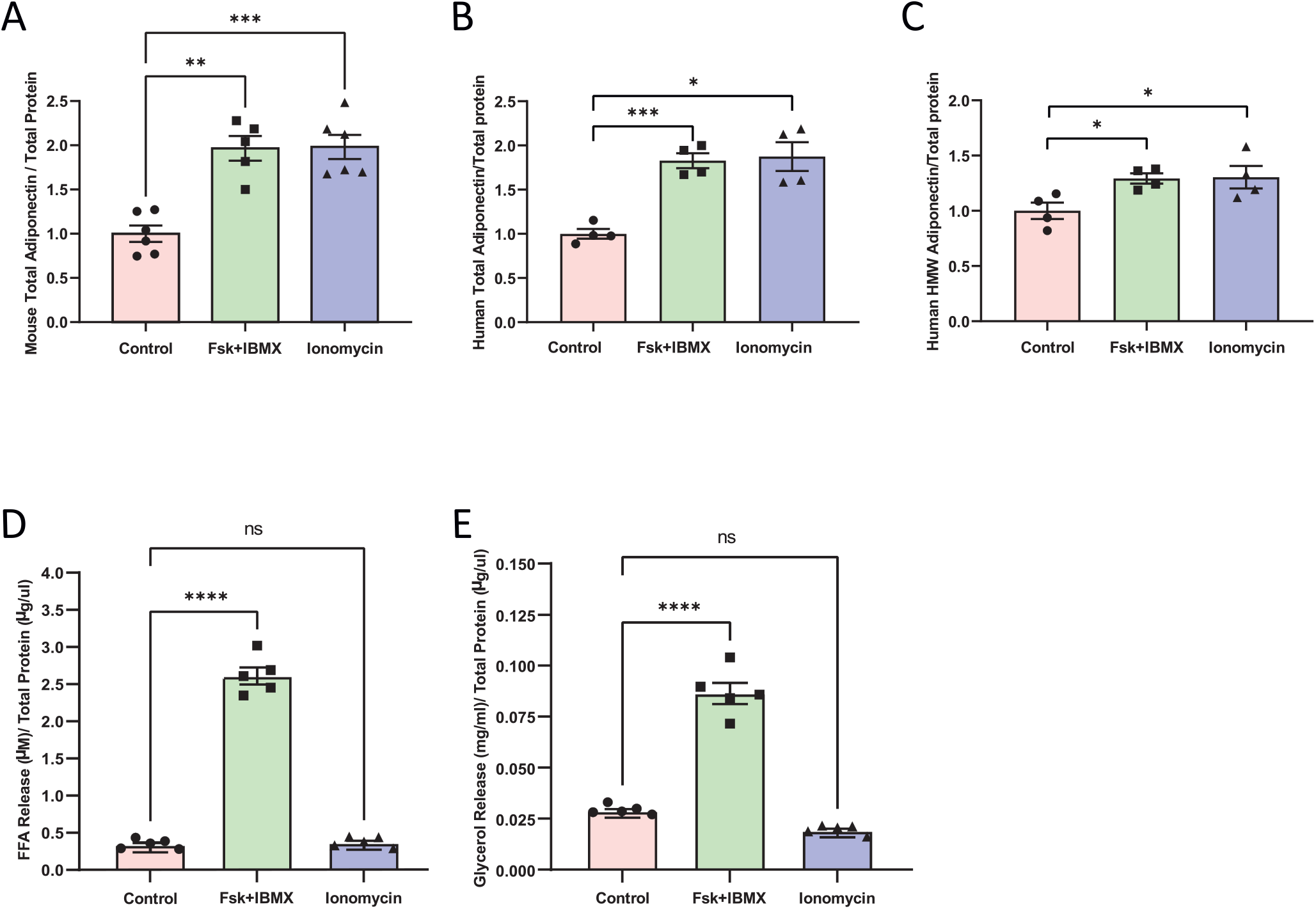
cAMP- and Ca2+-stimulated adiponectin secretion in native and adiponectin-mCherry-expressing 3T3-L1 adipocytes. **A)** Native mature 3T3-L1 adipocytes were serum starved for 30 min at 32^0^C and treated with DMSO, FSK/IBMX, or ionomycin for 30 min at 32^0^C. The extracellular solutions were collected and analysed for endogenous total adiponectin release using ELISA. Statistical analysis: Brown-Forsythe and Welch ANOVA (Dunnett’s T3 multiple comparisons test), assuming unequal SDs at α – level of 0.05. N=6 for Control and ionomycin; N= 5 for FSK/IBMX. **B)** Human total adiponectin and **C)** human HMW adiponectin secretion in stably expressing adipocytes, measurement using ELISA. Statistical analysis: Ordinary One-way ANOVA (Holm-Sidak’s multiple comparisons test), assuming equal SDs at α – level of 0.05. N= 4. Data are presented as mean ± SEM. *P<0.05, **P<0.01 and ***P<0.001. Lipolysis measured as FFA **D)** and glycerol **E)** release in 3T3-L1 adipocytes stably expressing recombinant adiponectin under DMSO (Control), FSK/IBMX or ionomycin stimulation. Note that only co-stimulation with FSK/IBMX significantly increased the secretion of FFA and glycerol compared to the control. Statistical analysis: Ordinary One-way ANOVA (Dunnett’s multiple comparisons test), assuming equal SDs at α – level of 0.05. N=5. Data are presented as mean ± SEM. ****P<0.0001 and ns = non-significant.

## Discussion

Pre-stored neurotransmitter- or hormone-containing vesicles typically release their content to the extracellular space upon receiving the correct triggering signal. Classically, an elevation of cytosolic Ca^2+^, released from intracellular stores or entering via Ca^2+^-permeable ion channels in the plasma membrane, stimulates vesicle exocytosis. Other intracellular mediators such as cAMP and ATP also have important roles for vesicle maturation steps (15). Peptide hormone exocytosis has been detailed in a number of endocrine cell types, including insulin secreting beta-cells (16) and catecholamine-releasing chromaffin cells (17). However, the control of hormone release in white adipocytes is a research field that has received little attention. Adiponectin, one of the chief hormones secreted from white adipocytes and the most abundant protein present in serum (18), has in several studies been reported to have favourable metabolic effects and to protect from development of obesity-associated metabolic diseases (1, 2). Although the general control of adiponectin secretion over shorter and longer time-periods has been addressed in a small number of studies (4, 19–21), adiponectin vesicle exocytosis – the molecular and cellular process in which adiponectin-containing vesicles are recruited to and fuse with the adipocyte plasma membrane to release their cargo - is a largely unexplored field. Here we have generated 3T3-L1 cells stably expressing human adiponectin fused with the fluorescent protein mCherry, and we use the overexpressing matured adipocytes to visualise and quantify adiponectin exocytosis. We demonstrate that adiponectin vesicle trafficking and exocytosis can be studied at the single vesicle level, and we determine that an elevation of intracellular Ca^2+^ stimulates adiponectin vesicles exocytosis in the absence of a concomitant increase of cAMP. Below we discuss the usefulness of our cells for future studies of adiponectin vesicle behaviour, and we reconsider the Ca^2+^-dependence of adiponectin exocytosis.

### The adiponectin-mCherry protein is targeted to endogenous adiponectin vesicles that resemble hormone-containing vesicles in other endocrine cell types

The addition of a fluorescent tag to a protein may disturb the correct targeting of the product. Our TIRF data propose that the chimeric protein is present in vesicular compartments, indicated by the punctate appearance of the fluorescence signal. Moreover, the real-time recordings suggest that the overexpressing adipocytes are functional, and that adiponectin-mCherry is targeted to the correct vesicle population; this is shown by that the labelled vesicles fuse with the plasma membrane in response to signals known to stimulate adiponectin exocytosis. In further support of this, measurements of released human adiponectin show that the fusion protein is secreted in response to an elevation of cAMP or Ca^2+^. Our estimated diameter of 200-300 nm for the adiponectin-labelled vesicles is similar to hormone-containing vesicles in other endocrine cell types, such as insulin granules with a diameter of ∼300 nm (26).

We have previously demonstrated that secretion of the hormones leptin, apelin and adipsin is unaffected by cAMP in 3T3-L1 adipocytes (5) and that resistin is co-secreted with adiponectin (contained within the same vesicles (37)). Collectively, our data propose that the rapid loss of fluorescence puncta in response to cAMP or Ca^2+^ reflects exocytosis of single vesicles belonging to the endogenous adiponectin vesicle population.

### Both cAMP and Ca^2+^ trigger exocytosis of adiponectin-mCherry labelled vesicles

The real-time imaging data in Figs. 4 and 5, as well as Suppl. videos 1 and 3, confirm the previously shown importance of cAMP and Ca^2+^ for stimulation of adiponectin vesicle release (5–7, 27). However, the results in Fig. 5 and Suppl. video 3, where ionomycin triggers exocytosis in the absence of a cAMP increase, can appear to contradict previous experiments showing that Ca^2+^ alone is unable to induce exocytosis in 3T3-L1 adipocytes (6, 7). Notably, there are some important difference between our previous patch-clamp data and the current TIRF recordings: During the electrophysiological experiments, the interior of the adipocyte is controlled by the pipette solution – thus by exclusion of cAMP in the pipette solution washing into the cell, the experimental approach applied in (6, 7), the 3T3-L1 adipocytes can essentially be depleted of cAMP. In contrast, the adipocytes used for TIRF imaging are metabolically intact and have a continuous endogenous production of cAMP, maintaining basal levels of the cyclic nucleotide. We suggest that the elevated Ca^2+^ acts together with preserved endogenous levels of cAMP, to stimulate adiponectin exocytosis in micro-domains where cAMP is sufficiently high. The finding that ionomycin on its own stimulates adiponectin secretion in metabolically intact adipocytes is in agreement with data on secretion of the adipocyte hormone resistin (28); we recently showed that resistin is co-secreted with adiponectin in mouse adipocytes (thus contained within the same vesicles) and that ionomycin induced the release of resistin in the absence of a cAMP-elevating agent (29). An alternative explanation to the effect of ionomycin alone is that the increase in intracellular Ca^2+^ activates Ca^2+^-dependent adenylyl cyclases leading to production of cAMP. In Musovic et al. 2021, we were unable to demonstrate a significant increase of cAMP levels in 3T3-L1 adipocytes exposed to ionomycin (Fig. 1C of (29)). However, cAMP was measured in cell lysates, with no information of changes of cAMP confined to specific near-membrane regions. Thus, ionomycin might stimulate local elevations of cAMP, at sites of adiponectin vesicle exocytosis. It is well known that cAMP is compartmentalised in adipocytes and that this regulates several intracellular processes (30). A role of caveolae for the spatial control of cAMP has been demonstrated in cardiac myocytes, and proposed to be relevant for βAR signalling (31). We have previously shown that caveolae are necessary for the organisation of signalling pathways involved in the regulation of HMW adiponectin exocytosis (32).

Vesicles that contain peptide/protein hormones typically undergo a series of maturation steps including physical movement, docking (stable attachment to the plasma membrane) and priming (chemical modifications that make docked vesicles release competent) before they can discharge their cargo upon fusion with the plasma membrane (15). The data in Figs. 4 and 5 as well as supplementary videos 1 and 3, show that both cAMP and Ca^2+^ trigger exocytosis of adiponectin vesicles that are already docked at the adipocyte plasma membrane; no new vesicles (newcomers) are observed to enter the TIRF zone. This observation seems to stand in conflict with our previous electrophysiological data demonstrating that an elevation of Ca^2+^ maintains adiponectin secretion over time, and with our conclusion that Ca^2+^ induces vesicle replenishment. In this context, it is essential to keep in mind that we image only a small part of/near the adipocyte plasma membrane, the ventral side of the cell attached to the culture dish, whereas the capacitance recording reflect exocytosis that occur over the entire cell membrane. Consequently, vesicles that move to and undergo exocytosis at the apical side of the adipocyte, outside of the TIRF imaging zone, are not visualised.

### Choice of cells and fluorescent protein

3T3-L1 cells were used to generate the adiponectin-mCherry expressing adipocytes. A number of studies have demonstrated that *in vitro* differentiated 3T3-L1 adipocytes are a relevant cell model for investigations of the adiponectin secretion process (5–7, 9, 22). In fact, all mechanisms and mediators determined to regulate adiponectin exocytosis in 3T3-L1 adipocytes have been confirmed in primary mouse and human adipocytes ((7–9) and Musovic & Olofsson, unpublished data). The 3T3-L1 cells are easy to culture and differentiate, and their functionality with regard to adipocyte metabolic processes, such as lipolysis (23) and glucose uptake (24), is well determined. Nonetheless, they are clonal cells and we acknowledge that results must be continuously validated in experiments with primary, preferably human, adipocytes. A disadvantage with the 3T3-L1 adipocytes is that their ability to differentiate into mature adipocytes ceases with increased cell passage. Although multiple vials of low passage cells can be stored, it will eventually be necessary to produce new stably expressing 3T3-L1 cells. However, we are not aware of any adipocyte cell line that does not have this limitation. Collectively, we regard the 3T3-L1 adipocytes to be a good choice for our studies.

A large number of fluorescent proteins are available for protein tagging and real-time studies of their specific function. The continuous development of fluorescent proteins with enhanced or altered qualities have yielded a supply that covers the full visible spectrum, with diverse advantages and disadvantages. Introduced nearly a decade ago, mCherry remains a frequently used fluorescent marker for cellular studies. The mCherry has several advantages: It is bright and it matures rapidly, enabling experiments to be performed soon after initiation of expression. It is also highly photostable and resistant to photobleaching (25). Since experiments must be carried out at high time-resolution and over an extended time-period (up to 30 min) the listed qualities are important. We therefore labelled adiponectin with mCherry, expressed under the CMV promoter that has high housekeeping expression. Human adiponectin was chosen to enable us to biochemically distinguish between endogenous mouse adiponectin and the overexpressed fusion protein. Nonetheless, there are some disadvantages related to using mCherry for studies of adiponectin exocytosis: Red fluorescent proteins display limited pH sensitivity compared to e.g. enhanced green fluorescent protein (EGFP) or pHluorin. Upon vesicle fusion with the plasma membrane the pH of the vesicle lumen (which is acidic) increases; pH-sensitive fluorescent proteins will consequently increase in brightness already upon formation of the vesicle-plasma membrane fusion pore, thus confirming vesicle fusion.

For this reason, we initially evaluated the use of EGFP for studies of adiponectin exocytosis, but both the more rapid photobleaching of EGFP, and lipid autofluorescence interfering with the green signal complicated the studies. With regard to pHluorin, this optical indicator is non-fluorescent at low pH and thus does now allow investigations of events that occur prior to vesicle fusion with the plasma membrane. We nonetheless acknowledge the advantage of using a pH-sensitive fluorescent tag for studies of vesicle fusion and cargo release and we will evaluate their use for future investigations of the adiponectin exocytosis process. We also recognise that tagging adiponectin itself might interfere with both protein mulitmerisation and the secretion process. However, the secretion data confirm that the stably expressing adipocytes release abundant amounts of human HMW adiponectin, verifying multimerisation of the chimeric protein. Regardless of this, it would be valuable to assess if labelling of another exocytosis vesicle marker such as Neuropeptide Y (NPY), a proven reliable marker for insulin granule exocytosis (25), is useful for real-time imaging of the adiponectin vesicle release process.

### What can we learn about the control of adiponectin secretion by real-time TIRF imaging of adiponectin exocytosis?

Our previous work has begun to detail the white adipocyte adiponectin exocytosis process (5–9, 22, 27, 33). However, the combination of methods used have temporal and spatial limitations and detailed knowledge of adiponectin vesicle dynamics and release can only be gained by high-resolution imaging in live cells. The finding that adiponectin exocytosis is similarly controlled in 3T3-L1 and primary mouse and human adipocytes (5, 7–9, 32, 33) open opportunities to use the stably expressing cells to achieve a comprehensive understanding of mechanisms and mediators that control adiponectin vesicle exocytosis, at the cellular and molecular level. TIRF movies can be acquired at high time resolution (10-20 FPS), thus allowing real time direct visualisation of the adiponectin vesicle exocytosis process. Consequently, the control and rate of docking and exocytosis can be determined.

In summary, TIRF imaging of cultured or primary live adipocytes with fluorescently labelled adiponectin vesicles will enable us, and others, to define currently unknown regulatory steps of adiponectin vesicle dynamic, maturation and exocytosis, at a level of detail that has not been studied previously. How are adiponectin vesicles recruited to the exocytosis site? Are the vesicles present before, during and after exocytosis? Do vesicles interact with each other during the exocytosis process? Those are questions that can be answered with the type of research presented here.

## Author contributions

MMS and CSO outlined the study and designed the experiments. MMS carried out all experimental work and analysis. CSO and MMS wrote the manuscript. SB provided method and analysis expertise and gave valuable input on the manuscript. All authors approved the final version of the manuscript.

## Abbreviation

AAV: Adeno-Associated Virus
ANOVA: Analysis of variance
ATCC: American Type Culture Collection
cAMP: Cyclic Adenosine Monophosphate
CMV: Cytomegalovirus
DMEM: Dulbecco’s Modified Eagle Medium
DMSO: Dimethyl sulfoxide
EDTA: Ethylenediaminetetraacetic acid
EGFP: Enhanced Green Fluorescent Protein
ELISA: Enzyme-linked immunosorbent assay
Epac1: Exchange factor directly activated by cAMP 1
Epac2: Exchange factor directly activated by cAMP 2
EC: Extracellular
FBS: Fetal bovine serum
FPS: Frames per second
FSK: Forskolin
GFP: Green Fluorescent Protein
HMW: High Molecular Weight
IBMX: 3-isobutyl-1-methylxanthine
NPY: Neuropeptide Y
NS: Not significant
PAGE: Polyacrylamide gel electrophoresis
PVDF: Polyvinylidene fluoride
qPCR: quantitative polymerase chain reaction
ROI: Region of interest
SD: Standard deviation
SDS: Sodium dodecyl-sulfate
SEM: Standard error of mean
shRNA: short hairpin ribonucleic acid
TIRF: Total internal reflection fluorescence

## Supplementary Videos

**Supplementary video 1**

3T3-L1 adipocytes stably expressing adiponectin-mCherry were treated with 10 µM Forskolin and 200 µM IBMX for 15 minutes during which images were taken at 2 FPS using TIRF microscopy. The video is condensed to show every 10^th^ frame. Time point of FSK/IBMX addition = 30 s. Scale bar = 10 µm.

**Supplementary video 2**

3T3-L1 adipocytes stably expressing recombinant adiponectin were treated with DMSO (control for Forskolin/IBMX recording) for 15 minutes during which images were taken at 2 FPS using TIRF microscope. The video is condensed to show every 10^th^ frame. Scale bar = 10 µm.

**Supplementary video 3**

3T3-L1 adipocytes stably expressing recombinant adiponectin were treated with 1 µM ionomycin for 15 minutes during which images were taken at 2 FPS using TIRF microscope. The video is condensed to show every 10^th^ frame. Time point of ionomycin addition = 30 s. Scale bar = 10 µm.

**Supplementary video 4**

3T3-L1 adipocytes stably expressing recombinant adiponectin were treated with DMSO (control for ionomycin recording) for 15 minutes during which images were taken at 2 FPS using TIRF microscope. The video is condensed to show every 10^th^ frame. Scale bar = 10 µm.

---

**Supplementary Fig 1:**
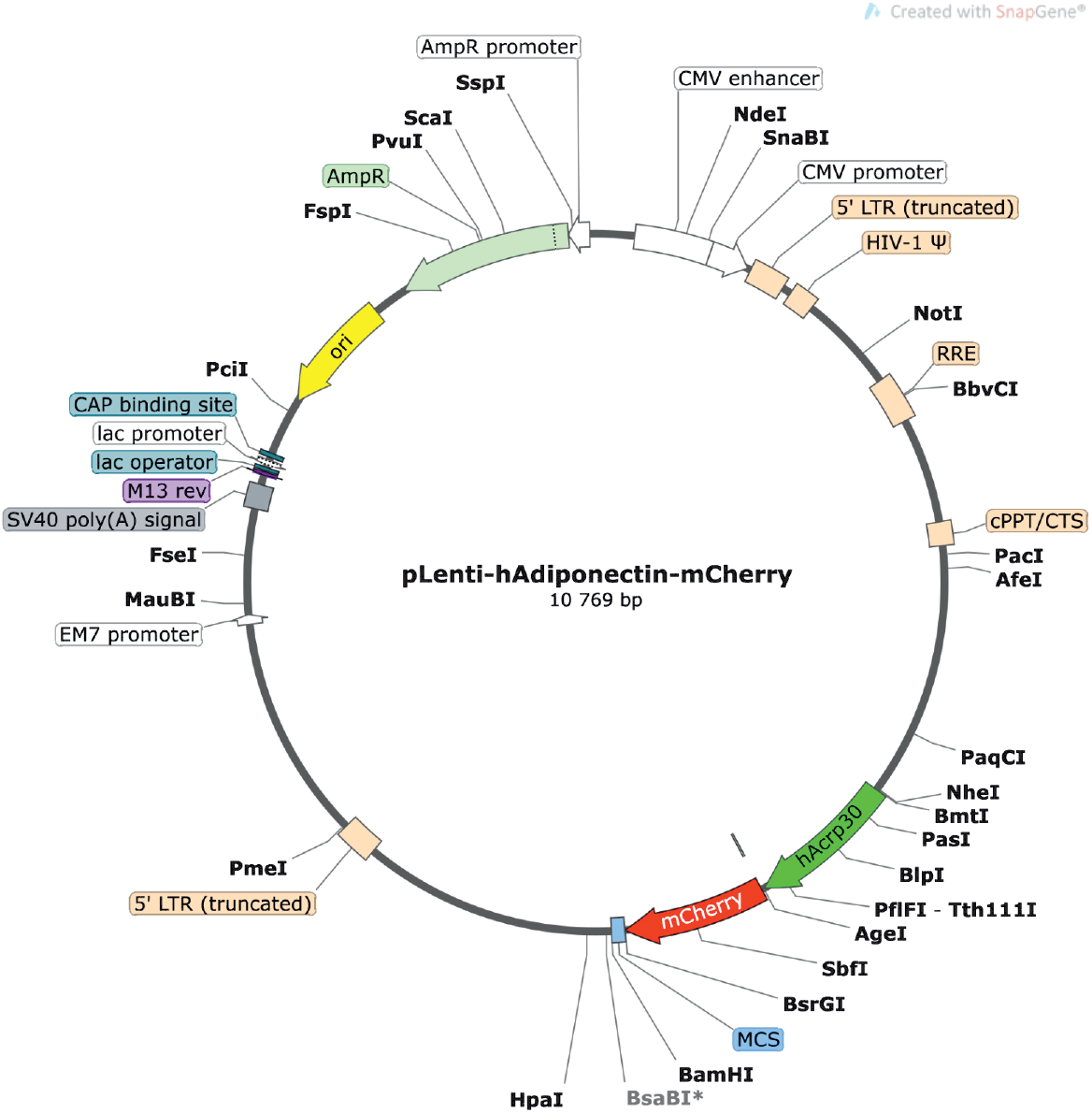
Plasmid map of lenti-viral construct for human adiponectin fused with mCherry at C-terminal created with SnapGene. 3T3-L1 adipocytes expressing recombinant adiponectin were serum starved for 30 minutes at 32 ^0^C and treated with 10 µM Forskolin and 200 µM IBMX, 1 µM ionomycin and DMSO as Control for 15 minutes at 32 ^0^C during which images were taken at 2 FPS using TIRF microscope.

